# Whole-brain EEG dynamics depending on the stimulus modality and task requirements in oddball tasks

**DOI:** 10.1101/2025.04.15.648394

**Authors:** Shiho Kashihara, Tomohisa Asai, Hiroshi Imamizu

## Abstract

Electroencephalography (EEG) microstates constitute temporal map configurations that reflect the whole brain electrical state. The dynamics of EEG microstates may serve as an effective discretization method for capturing spatiotemporally continuous neural dynamics with high temporal resolution. In this study, we employed polarity-sensitive microstate analysis to investigate whole-brain state transitions during audiovisual oddball tasks. Moreover, we examined how sensory modality and its coupling, types of response to target stimuli, and the physical presence or absence of target stimuli affected EEG dynamics. The results demonstrated that the abovementioned factors affected both behavioral indices and the event-related potential (ERP) components, particularly the P300. Importantly, when considering the topographical polarity of map configuration, transitions to microstate E-, which originates within 300 to 600 ms after stimulus onset and coincides with the typical latency of the P300 component potentially reflect attentional and conscious processes that are associated with the P300. These novel insights into the dynamic transitions of whole brain states during cognitive processes complement the results of traditional ERP analyses.

## 1. <u>Introduction</u>

Cognitive functions, such as recognizing and responding to external stimuli, are mediated by neural activity in the brain. Traditional approaches in cognitive neuroscience, known as the “Sherringtonian view,” attributed specific cognitive functions to localized brain areas and circuits (Barack & Krakauer, 2021). However, the “Hopfieldian view” or “population doctrine” constitutes a paradigm shift wherein whole-brain dynamics and neural state-space transformations explain cognition (Barack & Krakauer, 2021; Ebitz & Hayden, 2021). Studies using functional magnetic resonance imaging (fMRI) and intracranial electrodes described cognition in terms of neural state-space transitions (Iyer et al., 2022; Mau et al., 2020; Pessoa, 2019; Shine et al., 2019); however, noninvasive methods with high temporal resolution, such as scalp EEG, offer promising alternative approaches for investigating the neural dynamics that underlie cognition.

EEG microstate analysis provides a promising method for capturing whole-brain neural activity with high temporal resolution (e.g., Khanna et al., 2015; D. Lehmann et al., 1987; Michel & Koenig, 2018; Mishra et al., 2020). EEG microstates are quasi-stable electric field patterns in scalp EEG that reflect the globally synchronized activity of large neural networks (D. Lehmann et al., 1987; Michel & Koenig, 2018). Frequently referred to as the “atoms of thought” (Dietrich Lehmann, 1990), EEG microstates have been extensively studied in relation to psychiatric disorders, consciousness, and cognition.

To explore the functional role of EEG microstates, studies that used simultaneous EEG-fMRI and EEG source localization linked specific microstates to distinct brain networks (Al Zoubi et al., 2022; Britz et al., 2010; Musso et al., 2010; Yuan et al., 2012). Furthermore, researchers have investigated whether particular microstates emerge during cognitive tasks and how task characteristics influence microstate dynamics (Bréchet et al., 2019; D’Croz-Baron et al., 2021; Kim et al., 2021; Seitzman et al., 2017). For instance, Milz et al. (2016) found that microstate A and B (msA and msB) parameters increased during visualization and verbalization tasks, respectively. More recently, D’Croz-Baron et al. (2021) analyzed EEG topographical maps during auditory and visual encoding and recognition tasks, and reported the increased occurrence of msB following visual stimulus encoding. However, the relationship between specific microstates and cognitive functions remains inconsistent and is sometimes contradictory, which leaves their precise role unclear.

Previous studies have primarily focused on describing conditional differences in the discrete, static properties of EEG microstates, such as their frequency of occurrence and duration. However, recent research has challenged the discrete nature of microstates and emphasized the importance of considering the continuous dynamics that underlie EEG microstate sequences (Asai et al., 2023; Kashihara et al., in press; Michel & Koenig, 2018; Mishra et al., 2020; Shaw et al., 2019; Sikka et al., 2020; Van de Ville et al., 2010). Despite the growing body of knowledge on discrete microstate properties and the recognition of their continuous dynamics, the dynamic patterns of EEG microstate transition that emerge in response to external stimuli remain poorly understood. In contrast, Tamano et al. (2022) reported microstate dynamics during an N-back task and identified the relationship of certain transition probabilities comprising polarity with performance on the N-back task. However, these authors did not consider topographical polarity a better discretization method, assuming the continuity of EEG dynamics, and the significance of polarity consideration was limited. In this study, we address the knowledge gap indicated by this issue by using a polarity-sensitive microstate analysis and focusing on their transitions, while considering the continuous trajectory of the essentially continuous EEG dynamics in the state space (Kashihara et al., in press).

This study aimed to investigate EEG dynamics during cognitive tasks from the perspective of whole-brain state transitions while considering the spatiotemporal continuity of EEG activity. Instead of the traditional Sherringtonian approach of event-related potentials (ERPs), we sought to elucidate the neural basis of cognition through whole-brain state transition dynamics in the neural state space. To achieve this, we introduced a polarity-sensitive microstate analysis (Kashihara et al., in press) to descriptively analyze the stimulus-locked whole-brain state dynamics. Specifically, we extended the conventional ERP microstate approach (Kindler et al., 2011; Kondakor et al., 1997; D. Lehmann et al., 1994; Pizzagalli et al., 2000; Ruggeri et al., 2019) by incorporating topographical polarity, and thereby enabled a more precise characterization of state transitions in EEG activity. Previous studies on ERP microstates have examined how pre-stimulus topographical configurations influenced responses in oddball tasks (Kondakor et al., 1997; D. Lehmann et al., 1994). Research conducted in recent years has elucidated EEG microstate topography sequences that reflect the conflict-monitoring processes during the Stroop task (Ruggeri et al., 2019) and EEG microstate dynamics that predict working memory performance during the N-back tasks (Tamano et al., 2022). By incorporating topographical polarity, our approach enables the appropriate discretization of stimulus-locked, inherently continuous spatiotemporal EEG transitions while preserving trajectory alignment in the neural state-space based on topographical dissimilarity. This refinement enhances the ability to more accurately capture the dynamics of overlapping cognitive processes.

Therefore, we decided to use an oddball task as a cognitive task. The oddball paradigm, in which two or more stimulus events are presented at different frequencies, is a well-known procedure in ERP studies because it allows the measurement of neural processes related to stimulus perception and internal processing. In particular, among the salient ERP components that are elicited by this task, P300 – a peak that appears in the positive direction approximately 300 ms after target-stimulus presentation – has been used to retrieve endogenous mental activity. Furthermore, P300 may represent brain activity during the modification of a model of the environment, that is, during a contextual update (Donchin & Coles, 1988). Thus, by using the oddball task, wherein the association of ERP responses and cognitive processes has been well examined, clues to examine the functional role of EEG microstate dynamics may be found.

Taken together, this study aims to clarify the functional role of continuous whole-brain state dynamics during task performance by designing the task features of the oddball tasks in accordance with the findings accumulated in previous ERP studies. Therefore, we conducted experiments wherein several factors that influence the P300 component were manipulated to examine differences in microstate responses. Specifically, the following three points were investigated: the influence of the sensory modality of the stimulus (Experiment 1), the influence of the response format to the target (Experiment 2), and the influence of the presence or absence of physical stimuli on the microstate dynamics of the target (Experiment 3).

First, as a factor that affects the P300, the sensory modality of the stimulus (auditory or visual) affects the amplitude and latency of P300 (Polich & Heine, 1996). Furthermore, when the standard and target stimuli are intermodal (e.g., visual standard and auditory target), the late ERP component, including P3, increased, as compared to that in the unimodal case (Brown et al., 2006, 2007). Considering the possibility that sensory modality-specific processing may be associated with specific classes of EEG microstates (e.g., Britz et al., 2010), it is possible that the effects of the stimulus modality itself and its coupling with stimulus conditions can also be ascertained in the emerged EEG microstate classes or dynamics even when examining continuous whole-brain state transitions. Therefore, we prepared oddball tasks with different stimulus modalities and coupled them with conditions (unimodal auditory task, i.e., auditory target–auditory standard; unimodal visual task, i.e., visual target–visual standard; inter-modal auditory-target task, that is, the auditory target–visual standard; intermodal visual-target task, that is, visual target–auditory standard). First, we examined the effects of the modalities and their coupling on EEG microstate dynamics (in Experiment 1).

Second, we examined the effect of the response format required for the target stimulus. In the context of ERP studies, the P3 amplitude or latency may differ depending on the response format, such as whether the target stimulus requires a key press response or mental count. The findings pertaining to the effect of the response mode on ERPs are confusing. Some studies reported no difference between key-press and count for the target stimuli (Starr et al., 1995), whereas others reported that the P3 in the key-press task was greater than that in the counting task (Brázdil et al., 2003; Kotchoubey, 2014); moreover, there were even contradictory results (Barrett et al., 1987; Salisbury et al., 2001). As some findings suggest that motor behavior affects the discrete parameters of the EEG microstate (Croce et al., 2022), we examined the effects of the response format on EEG microstate dynamics by establishing a count-response group that can isolate the influence of motor behavior compared to key presses (Experiment 1 vs. Experiment 2).

Third, we examined the effect of the presence/absence of physical target stimuli. P3 potentially reflects an endogenous component and appears even when the target stimulus is not physically present (Ragazzoni et al., 2019). By setting up a task that requires a key press when a regularly presented physical stimulus is missing (the omitted-target oddball task), we can isolate the influence of the physical properties of the target stimulus and examine how purely endogenous processes of the target can be involved in continuous whole-brain state transitions (Experiment 1 vs. Experiment 3).

## 2. Methods

### 2-1. Participants

Three experiments involved 66 participants, and there was no duplication of participants. All experiments were approved by the ethics committee of the Advanced Telecommunications Research Institute International, Kyoto, Japan (approval numbers 21-144 and 21-143). All participants provided written informed consent before the experiment.

#### 2-1-1. Experiment 1

Twenty participants (12 females, 8 males; age, mean [SD]: 30.7 [6.9] years) participated in Experiment 1 and declared that they were right-handed. As no previous study has ascertained the EEG microstate transition indices during the oddball task, we determined the number of participants under an expected moderate effect size (Cohen’s *d* = 0.45) based on the robustness of the neural activity elicited by the oddball task (e.g., P300 ERP component). Using PANGEA v0.2 (https://jakewestfall.shinyapps.io/pangea/)(Power ANalysis for GEneral Anova designs, Westfall, 2016), we calculated the sample size required for detecting the interaction between stimulus modality and condition when assuming the experimental design of 2 (modality coupling: unimodal, inter-modal) ×2 (target stimulus modality: auditory, visual) ×2 (stimulus condition: target, standard) with the factor of participants as a random effect. We considered the repetition of stimulus presentations 80 times (in the target condition). Accordingly, PANGEA suggested 15 participants for power (1-β) > 0.80 with Cohen’s *d* = 0.45. We recruited 20 participants, considering the possibility of missing data.

#### 2-1-2. Experiment 2

The number of participants in Experiment 2 was determined according to the sample size in Experiment 1. Twenty-one participants (11 females and 10 males; age, mean [SD]: 24.8 [5.3] years) participated in Experiment 2. All participants were right-handed and had never participated in Experiments 1 and 3. One participant was excluded because of refusal to participate; therefore, data from 20 participants (10 females and 10 males; age, mean [SD]: 25.0 [5.3] years) were included in the analysis.

#### 2-1-3. Experiment 3

In total, 21 participants (11 males and 10 females; age, mean [SD]: 22.6 [4.2] years) were included in Experiment 3, were all right-handed, and had never participated in Experiments 1 and 3. Two participants were excluded because one refused to participate and the other experienced recording failure. Therefore, data from 19 participants (nine males and 10 females; age, mean [SD]: 22.9 [4.4] years) were used in the subsequent analyses.

### 2-2. Apparatus

Stimuli were presented on a 24-inch color LCD monitor in all three experiments (FlexScan EV 2456, EIZO, Japan), and responses were recorded via a computer keyboard in Experiments 1 and 3 (Realforce91U, Topre Corporation, Japan). The monitor (resolution: 1920 × 1200) was situated approximately 55 cm in front of the seated participants. The size of visual stimuli was approximately 480x480 pixels. Stimulus presentation and response collection were controlled by PsychoPy v2021.1.4 (J. Peirce et al., 2019; J. W. Peirce, 2007, 2008).

### 2-3. Tasks and materials

We conducted audiovisual oddball tasks in the three experiments with different task formats. In the task, two types of stimuli with different frequencies of the presentation were shown sequentially (high-frequency: standard, low-frequency: target). Stimuli were presented randomly, with the restriction that the target stimuli were not presented consecutively. Participants were asked to respond to a low-frequency stimulus (target) as quickly and accurately as possible; that is, they were to press the space key when the target is presented (Experiment 1), keep counting the number of target presentations in their mind (Experiment 2), and press the space key when the stimulus was absent (Experiment 3). In Experiment 3, only the standard stimulus was perceptible, and the target stimulus was a “missing target” that was physically absent. In general, a total of 400 trials were presented in each task. The rate of the presentation of the standard and target stimuli was 8:2. Therefore, the number of standard and target trials were 320 and 80, respectively. Each stimulus was presented for 200 ms, with a 1000-ms inter-stimulus interval.

#### 2-3-1. Experiments 1 and 2

In Experiments 1 and 2, there were four tasks based on the target stimulus modality (auditory or visual) and the modality coupling between target and standard stimulus (unimodal or inter-modal) as follows: unimodal-auditory oddball task, unimodal-visual oddball task, inter-modal auditory-target oddball task, and inter-modal visual-target oddball task (Figure 1). The order of tasks was partially counterbalanced; two unimodal tasks were conducted, followed by the inter-modal tasks, and the modality of the target stimulus was counterbalanced. Participants were asked to either press the space key (Experiment 1) or count the number of target stimuli in their minds (Experiment 2) when the target stimulus was presented.

**Figure 1.**
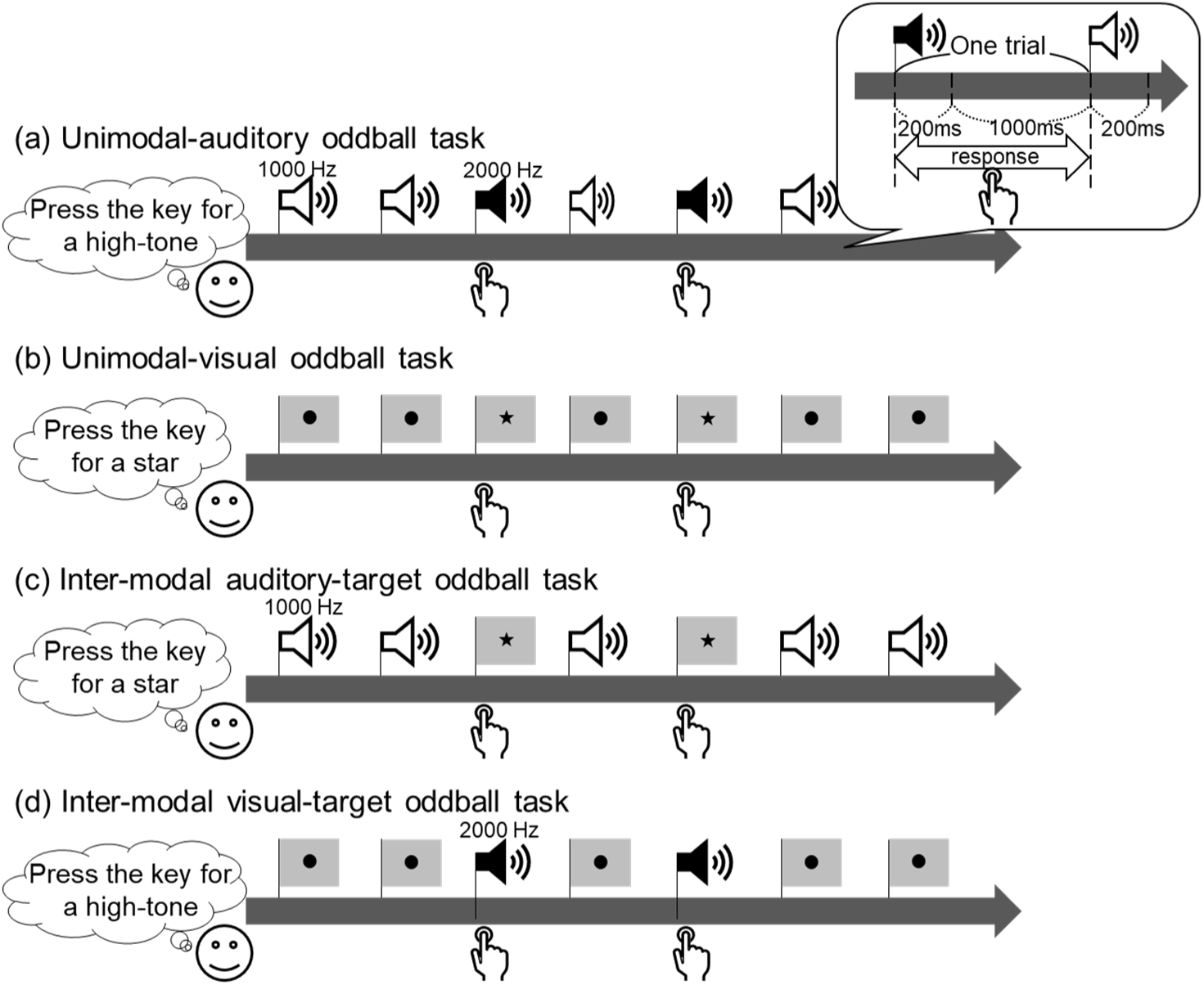
The overview of trials of each of the four types of oddball tasks. In each experiment, four different unimodal/inter-modal, auditory/visual tasks were performed with a combination of stimulus modalities (auditory or visual) for each stimulus with varying presentation frequencies (standard or target). In Experiment 3, only two unimodal tasks were performed because no physical stimulus was presented in the target trials. For each task, a total of 400 trials (320 standard and 80 target trials) of two blocks (200 trials, including 160 standard and 40 target trials) were presented. After finishing the first block, we asked the participants to take a break for an arbitrary time. Each stimulus was presented for 200 ms. The inter-stimulus interval was 1000ms. The order of stimulus presentation was determined pseudo-randomly for each participant under the condition that the target stimulus never appeared twice in a row. Participants were asked to press the space key as quickly and accurately as possible when the target stimulus was presented (Experiment 1), to count the number of times the target stimulus was presented (Experiment 2), or to press the space key as quickly and accurately as possible when they noticed a target trial where the stimulus was not presented (Experiment 3). We recorded the reaction time and correct/incorrect responses.

The experimental materials were 1000- and 2000-Hz pure tone for auditory stimuli, as well as a black circle (●) and a black star (★) for visual stimuli. The assignment of materials to stimulus conditions (i.e., standard or target condition) was fixed in Experiment 1 (auditory target: 2000 Hz, auditory standard: 1000 Hz, visual target: black star, visual standard: black circle). To offset the influence of materials, we counterbalanced the assignment of materials to conditions across participants in Experiment 2: for half of the participants (*N* = 10), the auditory target was a 2000-Hz tone, and the visual target was a black star; for the other half (*N* = 10), the auditory target was a 1000-Hz tone, and the visual target was a black circle.

#### 2-3-2. Experiment 3

The tasks in Experiment 3 were almost the same as those in Experiments 1 and 2. However, because the target stimulus was not physically present in Experiment 3 (“missing target”), there were only two unimodal tasks according to the standard stimulus modality (auditory or visual), and participants perceived only standard stimuli. The order of tasks was counterbalanced. In the tasks, auditory standard (1000 Hz) or visual standard stimuli (black circle) were presented at regular intervals, sometimes with missing stimuli (target; 20% of all trials). Participants were asked to press a space key when they noticed the missing target.

### 2-4. Procedure

The general procedure was common to all three experiments, except for what participants were to do during the oddball tasks. Participants wore an EEG cap, and the resting-state spontaneous EEG was recorded, for 5 minutes each, with eyes open and eyes closed. The participants sat in a comfortable chair and were asked not to think too deeply about certain things and to stay at rest—the order in which the eye-open and eye-closed resting recordings were counterbalanced. In eye-open resting EEG recording, the fixation cross appeared at the center of the monitor, and participants were asked to gaze at it. In the eyes-closed rest condition, participants were asked to keep their eyes closed throughout the recording and not to fall asleep.

Next, participants performed four oddball tasks; auditory or visual unimodal oddball tasks and inter-modal oddball tasks with auditory or visual targets. Inter-modal tasks were conducted after unimodal tasks, and the modalities were counterbalanced for both task presentation types. In one task, 400 trials were presented, with a break every 200 trials. The experiment lasted approximately 2 hours, from start to finish (Experiment 3 had two fewer tasks, but the time required for the experiment was the same as that for Experiments 1 and 2 because other tasks were performed that were not directly related to this study’s purpose).

Participants were given different instructions regarding task performance in each of the three experiments. In Experiment 1, participants were asked to press a space key as quickly and accurately as possible with either the index or middle finger of their right (i.e., dominant) hand when the target stimulus was presented. In Experiment 2, participants were asked to mentally count the number of target presentations and instructed not to speak aloud or use their fingers or bodies during counting. To increase engagement with the task, they were asked to answer the number of target presentations for that block every 200 trials. In Experiment 3, participants were given the same instruction as for Experiment 1; however, the target stimulus was not physically presented (“missing target”), and participants were asked to press the key when they noticed that stimuli presented at regular intervals in the target trials were missing.

### 2-5. EEG recording and preprocessing

EEGs were recorded in all three experiments using 32ch R-net (Brain Products GmbH, Gilching, Germany), silver–silver chloride electrodes attached to a saltwater sponge-based cap. A total of 32 electrodes were placed on the scalp based on the international 10-10 systems: that is, at Fp1, Fp2, Fz, F3, F4, F7, F8, F9, F10, FC1, FC2, FC5, FC6, Cz, C3, C4, T7, T8, CP1, CP2, CP5, CP6, Pz, P3, P4, P7, P8, P9, P10, Oz, O1, and O2. The ground and reference electrodes were placed at Fpz and FCz, respectively, and were off-line re-referenced to the average potential. The impedances were maintained under 50 kΩ. EEG signals were recorded at 500 Hz (Experiment 1) or 5000 Hz (Experiments 2 and 3) using an EEG amplifier (BrainAmp MR, Brain Products GmbH, Gilching, Germany) with an online bandpass of 0.016–250 Hz.

EEG data were preprocessed using an EEGLAB toolbox (EEGLAB v2019.1; Delorme & Makeig, 2004) on MATLAB R2019a (MathWorks). The raw EEG data were downsampled to 250 Hz and filtered using a finite impulse response (FIR) filter from 1 to 45 Hz. Next, we applied an EEGLAB plugin of *clean_rawdata* (v2.3) to the data to automatically identify bad channels and reconstruct bad segments using an artifact subspace reconstruction method (ASR; Mullen et al., 2015). The data was then run through adaptive mixture independent component analysis (AMICA) using an EEGLAB plugin of *AMICA* (v1.5.1) and also applied an EEGLAB plugin of *ICLabel* (v1.3; Pion-Tonachini et al., 2019) to identify artifact components. More specific information about the preprocess after filtering are as follow: First, the channel was rejected as a bad channel when it had a flat line of at least 5 s, had more than 4 *SD* line noise relative to its total channel signal, or correlated at less than 0.85 with adjacent channels. Second, continuous data were cleaned and reconstructed if their power deviated too much from the calibration data when evaluated by the ASR method. As the experimental booth was unshielded and the data were relatively noisy, we chose a 10-*SD* burst-detection criteria threshold as a relatively strict criterion (Chang et al., 2020). If the data that were decomposed into short segments had an RMS value greater than 10 *SD* compared to the clean-calibration data portion (the concatenated data was automatically determined based on the distribution of signal variance), the data in that time window were reconstructed based on the remaining components as containing artifacts. Third, we ran an adaptive mixture ICA (Hsu et al., 2018; J. A. Palmer et al., 2008; Jason A. Palmer et al., 2006) using an EEGLAB plugin of *AMICA*, and thereafter labeled each ICs using ICLabel (Pion-Tonachini et al., 2019) and conducted the dipole estimation using the EEGLAB plugin of *DIPFIT*. Finally, using this information, we kept the “signal IC”s that meet the following three criteria: (a) IC, which was labeled as “brain” by ICLabel; (b) IC in which the residual variance of the dipole was less than 15% (i.e., the IC with RV >15% is assumed to be mostly an artifact component); and (c) the dipole was presumed to be in the brain area. In addition, ICs that still contained data with amplitudes greater than 150 μV after these preprocess were rejected as likely to be artifactual components.

### 2-6. EEG analysis

#### 2-6-1. ERP analysis

Preprocessed EEG data were epoched from −200 to 1000 ms, with 0 ms at the stimulus onset. The trials that received incorrect responses were excluded from later analysis. Using these epochs, individual ERPs were computed for the two conditions: averaging target trials (M = 78.89, *SD* = 1.77; range: 68–80, for all four tasks) and averaging standard trials (M = 319.33, *SD* = 2.16; range: 303–320 for all of four tasks). Then, the individual ERPs were averaged into two grand-average ERPs for each condition. For the ERP analysis, the voltage was corrected using the segment from −200 to 0 ms as a baseline.

#### 2-6-2. EEG microstate analysis

We performed EEG microstate analysis on the preprocessed data using some of the functions of the EEGLAB plugin “*MST1.0*” (Poulsen et al., 2018). In the microstate analysis, the canonical microstate template is obtained through clustering topographies extracted from EEG or ERP signals (Khanna et al., 2014; D. Lehmann et al., 1998; Michel & Koenig, 2018; Poulsen et al., 2018). In general, in terms of the electric field stability (i.e., excellent signal-to-noise ratio), the points that receive clustering seeds are selected based on the peak of global field power (GFP; Khanna et al., 2014; D. Lehmann et al., 1998). After the templates are obtained, the topographic distribution at each timepoint in each participant’s EEG time series is labeled according to which class of template it most resembles (Khanna et al., 2014; D. Lehmann et al., 1998; Michel & Koenig, 2018; Poulsen et al., 2018).

In this study, we decided to use stable templates that were created in other large-scale EEG datasets (Kashihara et al., in press) as the templates for this study. This is because we considered microstates to be frequently traversed (“high traffic”) points that were defined in the standard MDS state space with embedded EEG topography (Asai et al., 2023; Kashihara et al., in press). Using this 10-state template that considers topographical polarity (Figure 2), we applied the template-fitting procedure to identify the label of the microstate class at each timepoint. The microstate template was fitted to individual trial-by-trial ERP epochs in the target and standard condition of each task.

**Figure 2.**
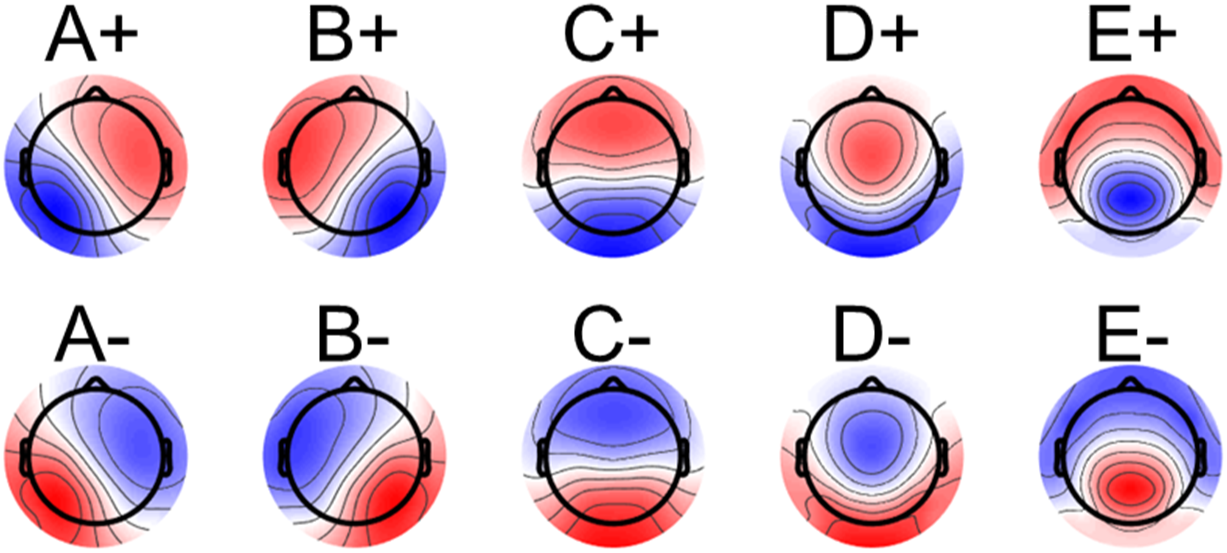
Stable EEG microstate templates considering topographical polarity from large-scale datasets. Microstate templates used for microstate fitting to EEG during oddball tasks in our study. These templates were originally created in a separate study by Kashihara et al. (in press) using a large-scale eyes-closed resting-state EEG dataset (N = 190, 8-min recordings). In their study, five microstate maps were initially extracted using the modified k-means method. To enable polarity-sensitive fitting, polarity-inverted patterns were then generated by reversing the sign of the voltage values at each electrode. Each state represents a stable topographical configuration of brain electrical activity. Considering topographical polarity facilitates an appropriate discretization of essentially spatiotemporally continuous EEG dynamics within the neural manifold space, where topographies are embedded using a multidimensional scaling method (Asai et al., 2023; Kashihara et al., in press).

## 3. Results

### 3-1. Behavioral results

#### 3-1-1. Experiment 1: Examining the effects of stimulus modality and coupling with conditions

We checked the mean correct rate for the key press of each task and found very high performance by the participants: 99.7% (*SD* = 0.3) for the unimodal-auditory task, 99.5% (*SD* = 0.9) for the unimodal-visual task, 99.8% (*SD* = 0.3) for the inter-modal auditory task, and 99.4% (*SD* = 1.6) for the inter-modal visual task. We conducted two-way ANOVA (target stimulus modality × modality coupling with stimulus condition) using the R packages of “anovakun (v4.8.9)” under R v4.4.2. Thus, there was neither significant main effects of the modality coupling with stimulus condition (*F*(1, 19) = 0.09, Greenhouse-Geisser’s ε = 0.39, *p* = .77, *η*_p_ ^2^ = .005) and the target stimulus modality (*F*(1, 19) = 1.40, Greenhouse-Geisser’s ε = 0.39, *p* = .25, *η*_p_ ^2^ = .07) nor an interaction effect for them (*F*(1, 19) = 1.42, Greenhouse-Geisser’s ε = 0.39, *p* = .25, *η*_p_ ^2^ = .07; Figure 3a).

**Figure 3.**
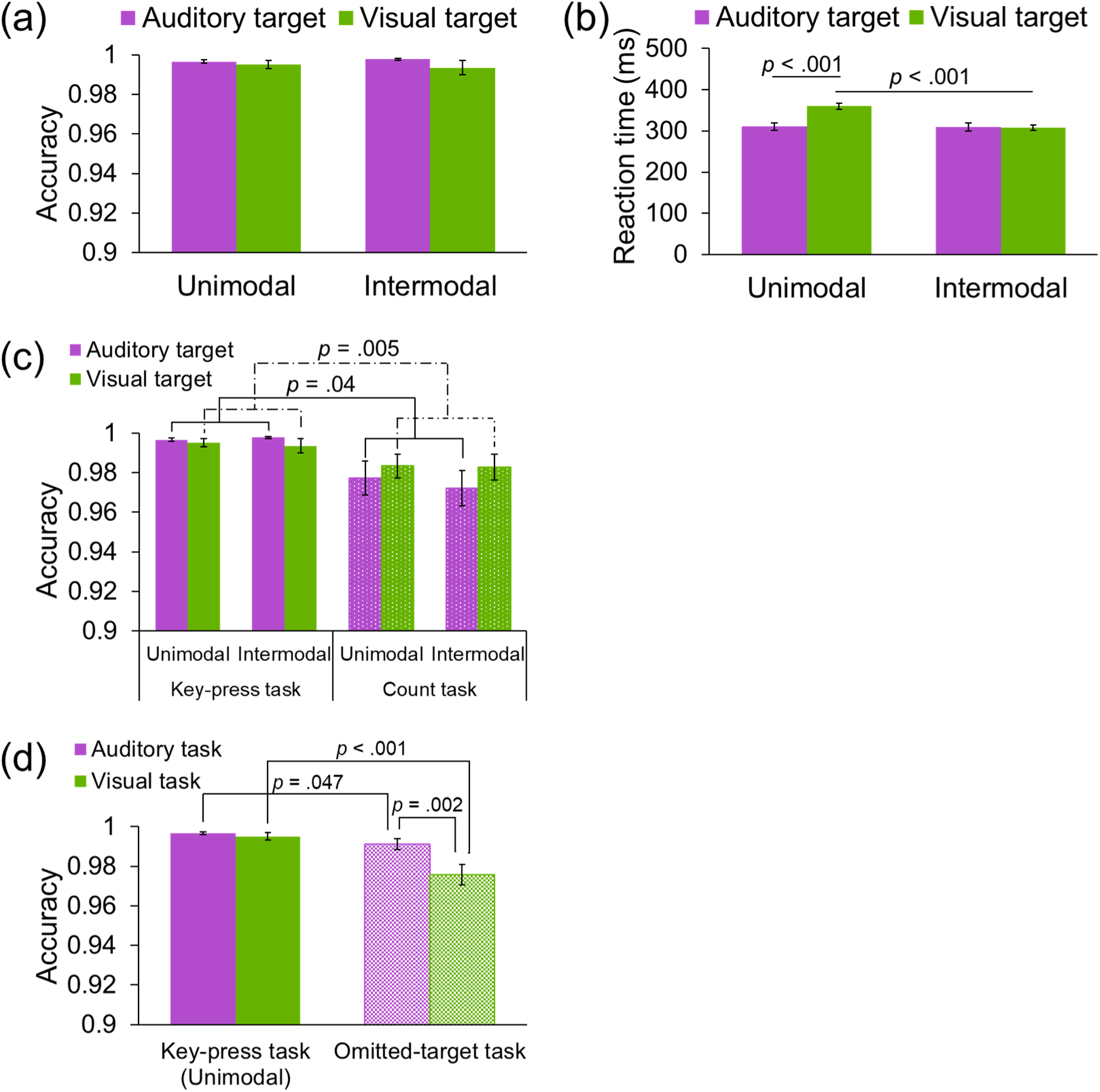
Behavioral results pertaining to the accuracy and reaction time in the three experiments in this study. Accuracy differences between target modality and modality coupling in Experiment 1. (b) Reaction-time differences between target modality and modality coupling in Experiment 1. (c) Accuracy differences between the target modality, modality coupling, and response format in Experiment 2. (d) Accuracy differences between stimulus modality and existence or absence of physical target stimuli in Experiment 3.

Furthermore, we checked the mean reaction time of each task in Experiment 1. The reaction times of all tasks were sufficiently fast: 311 ms (*SD* = 41) for the unimodal-auditory task, 360 ms (*SD* = 33) for the unimodal-visual task, 310 ms (*SD* = 45) for the inter-modal auditory task, and 308 ms (*SD* = 33) for the inter-modal visual task. We found significant main effects of the modality coupling with stimulus condition (unimodal > inter-modal: *F*(1,19) = 23.13, *p* < .001, *η*_p_ ^2^ = .55), the target stimulus modality (auditory < visual: *F*(1,19) = 19.24, *p* < .001, *η*_p_ ^2^ = .50), and their significant interaction effect (*F*(1,19) = 33.22, *p* < .001, *η*_p_ ^2^ = .64). The post analyses revealed significant differences between two visual-target tasks (unimodal visual > inter-modal visual-target: *F*(1,19) = 64.51, *p* < .001, *η*_p_ ^2^ = .77) and between two unimodal tasks (unimodal visual > unimodal auditory: *F*(1,19) = 39.82, *p* < .001, *η*_p_ ^2^ = .68; Figure 3b).

#### 3-1-2. Experiment 2: Examining the effects of the response format

We confirmed how correctly participants counted the targets for each task in Experiment 2. We calculated the counting accuracy for each task by absolutizing the magnitude of the discrepancy between the number of counts reported by participants (the total number of targets in two blocks of 200 trials) and the correct answer (80 targets) by dividing them by the total number. Even when the response format was changed to counting task, we found that the participants had very high performance: 97.7% (*SD* = 3.9) for the unimodal-auditory task, 98.3% (*SD* = 2.7) for the unimodal-visual task, 97.2% (*SD* = 4.0) for the inter-modal auditory task, and 98.3% (*SD* = 2.9) for the inter-modal visual task. A two-way ANOVA (target stimulus modality × modality coupling with stimulus condition) revealed that there was no significant effect of the modality coupling (*F*(1, 19) = 0.16, *p* = .69, *η*_p_ ^2^ = .009) and the target stimulus modality (*F*(1, 19) = 1.96, *p* = .18, *η*_p_ ^2^ = .09) and interaction between them (*F*(1, 19) = 0.16, *p* = .70, *η*_p_ ^2^ = .008) as in Experiment 1.

Next, we verified the effect of the response format on the behavioral index, that is, a comparison of the percentage of correct responses between Experiments 1 and 2. We conducted three-way mixed ANOVA (response mode × target stimulus modality × modality coupling with stimulus condition). Thus, there was a significant main effect of response mode (key press > count: *F*(1, 38) = 9.02, Greenhouse-Geisser’s ε = 0.92, *p* = .005, *η*_p_ ^2^ = .19). Furthermore, the primary interaction between the response mode and target stimulus modality was significant, which indicated that regardless of the modality coupling, when the target stimulus was visual and auditory, accuracy was higher in the key-press task than in the counting task, respectively (visual: *F*(1, 38) = 8.84, *p* = .005, *η*_p_ ^2^ = .19; auditory: *F*(1, 38) = 4.77, *p* = .04, *η*_p_ ^2^ = .11; Figure 3c). The main effects of other factors and their interactions were not significant (*p*s > .40).

#### 3-1-3. Experiment 3: Examining the effect of the presence or absence of physical **target stimuli**

The mean correct rates for the omitted targets were still high: 99.1% (*SD* = 1.1) for the auditory task and 97.6% (*SD* = 2.2) for the visual task. As shown with a paired *t*-test using the R function of *t.test* and package of “effsize (v0.8.1),” there was a significant difference between the two tasks (auditory > visual: *t*(18) = 3.64, *p* = .002, *d* = 0.80). In addition, the mean reaction time was 442 ms (*SD* = 55) in the auditory task and 567 ms (*SD* = 95) in the visual task. Moreover, we found a significant difference between the two tasks; participants had slower responses in the visual task than that in the auditory task (*t*(18) = 6.64, *p* < .001, *d* = 1.51).

In addition, we examined the effect of the physical presence of the target stimulus by comparing Experiments 1 and 3. Because Experiment 3 only included two types of tasks, one with auditory standard stimuli and one with visual standard stimuli, we decided to focus on the unimodal tasks in Experiment 1 and not consider the modality coupling of stimulus conditions here.

We conducted two-way mixed ANOVA (presence or absence of physical target stimulus × target stimulus modality) with accuracy as the dependent variable. Thus, there was a significant main effect of the presence or absence of physical target stimulus (existed target > omitted target: *F*(1, 37) = 12.10, Greenhouse-Geisser’s ε = 1.00, *p* = .001, *η*_p_ ^2^ = .25). In addition, there was a significant main effect of target stimulus modality (auditory > visual: *F*(1, 37) = 14.48, Greenhouse-Geisser’s ε = 1.00, *p* < .001, *η*_p_ ^2^ = .28). Furthermore, the interaction between these two factors was significant (*F*(1, 37) = 9.80, Greenhouse-Geisser’s ε = 1.00, *p* = .003, *η*_p_ ^2^ = .21). From the post analyses, there were significant differences between auditory tasks (existed target > omitted target: *F*(1,37) = 4.23, *p* = .047, *η*_p_ ^2^ = .10) and visual tasks (existed target > omitted target: *F*(1,37) = 13.27, *p* < .001, *η*_p_ ^2^ = .26). Moreover, in omitted target tasks, there were significant differences between standard stimulus modalities (auditory > visual: *F*(1,18) = 13.23, *p* = .002, *η*_p_^2^ = .42; Figure 3d).

### 3-2. ERPs

ERPs were calculated for each task and condition using 1200-ms epochs (−200 ms prestimulus to 1000-ms poststimulus). In Experiments 1 and 3, only correct trials (i.e., participants responded to the target stimulus and did not respond to the standard stimulus) were included in the later analysis.

#### 3-2-1. Confirming the P3 in each experiment: Comparison of standard and target **trials in each task of each experiment**

Figure 4 presents the grand average ERPs at a Pz electrode of each condition and task in Experiments 1–3. The results of the permutation test comparing the average ERPs for target trials and standard trials for each experiment and task showed that, in Experiment 1, the positive potential in the 300–400 ms range for target trials (magenta line) was significantly larger than that for standard trials (cyan) for each task, even after FDR correction (the following periods were significant at *p* < .05 after FDR correction for each task: unimodal-auditory, 264– 484, 532–540, and 928–936 ms; unimodal-visual, 92–112, 148–200, 304–516, 720–796, 804, 920–932, and 952–992 ms; inter-modal auditory-target, 180, 312–380 ms; inter-modal visual-target, 100–120, 164–200, 224–608, 684–696, and 988–1000 ms). This trend was almost same in Experiment 2 except for the inter-modal auditory-target task (unimodal-auditory: 332– 404, 436–484, 508–580, 588–616, 628–660, and 736–740 ms; unimodal-visual: 336–444 and 476–488 ms; inter-modal visual-target: -56, 24–28, 48–56, 92–96, 108–116, 172–196, 256–664, and 1000 ms; inter-modal visual-target: -60--56, 24-28, 48-56, 88-96, 108-116, 168-200, 252-620, 636-644, 1000 ms). Even in Experiment 3, where the target stimulus was not physically present, there was a significant difference between the target and standard trials in the auditory missing-target oddball task (−156 to −140, −28 to −8, 100–116, 148–164, 272– 616, 632–640, and 800–828 ms) as well as in the visual missing-target oddball task (136–188, 412–414, and 540–688 ms; *p*s < .05).

**Figure 4.**
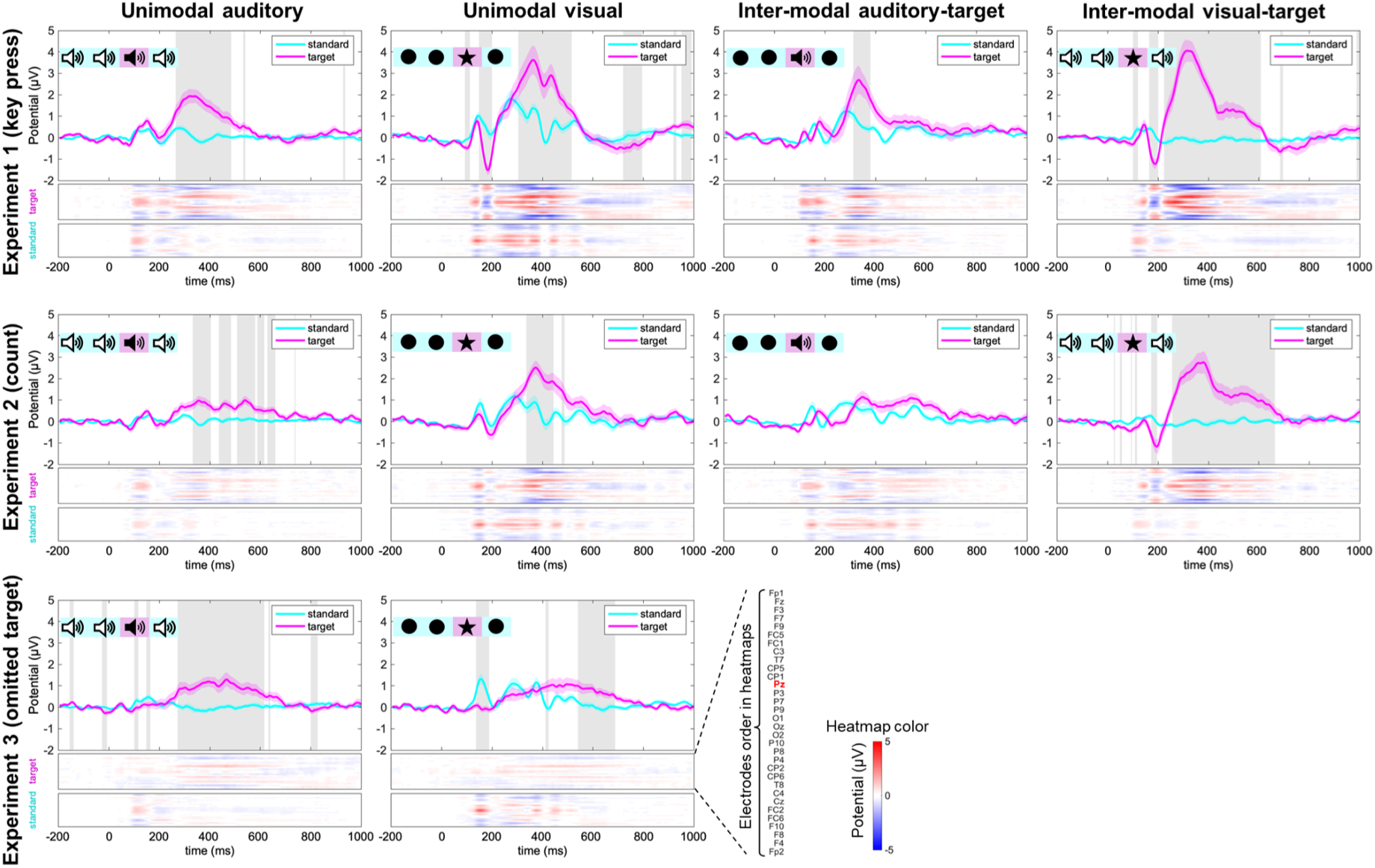
**ERPs at Pz and voltage heatmaps of all 32 channel electrodes of the target vs. standard trials for each task in three experiments.** The magenta (target trial) and cyan (standard trial) lines show the grand average ERP time course on the Pz electrode. Gray areas in each ERP panel mean a significant difference between the target and standard trial in t-tests with FDR correction. The heatmaps show the voltage level of all 32 electrodes. Of the two heatmaps in each task, the upper one depicts the grand average potential of the target trial, and the lower one is the standard trial. Red means positive potential whereas blue means negative potential (see the color bar). In each heatmap, the vertical axis shows the array of 32 electrodes, and the horizontal axis shows the time course. The order of the electrodes for all heatmaps is shown in the lower right corner.

#### 3-2-2. Experiment 1: Examining the effects of stimulus modality and coupling **with stimulus conditions**

Since the emergence of the P300 after the target stimulus was nearly confirmed, the peak amplitude at the Pz electrode between 300 and 600 ms in the trial average waveform of each participant was treated as the P300 amplitude in the subsequent ERP analysis.

First, to examine the effect of stimulus modality and modality coupling with stimulus condition, we performed a 2 (target modality: auditory, visual) × 2 (modality coupling: unimodal, intermodal) two-factor within-participant analysis of variance (ANOVA) for the mean P300 amplitude. Results showed that only the main effect of target modality was significant, with visual targets having larger P300 amplitudes than auditory targets (*F*(1,19) = 43.32, *p* < .001, η_p_ ^2^ = 0.55; Figure 5a). The main effect of modality coupling (*F*(1,19) = 2.43, *p* = .14, η_p_ ^2^ = .11) and interaction (*F*(1,19) = .07, *p* = .80, η_p_ ^2^ = .003) were not significant.

**Figure 5.**
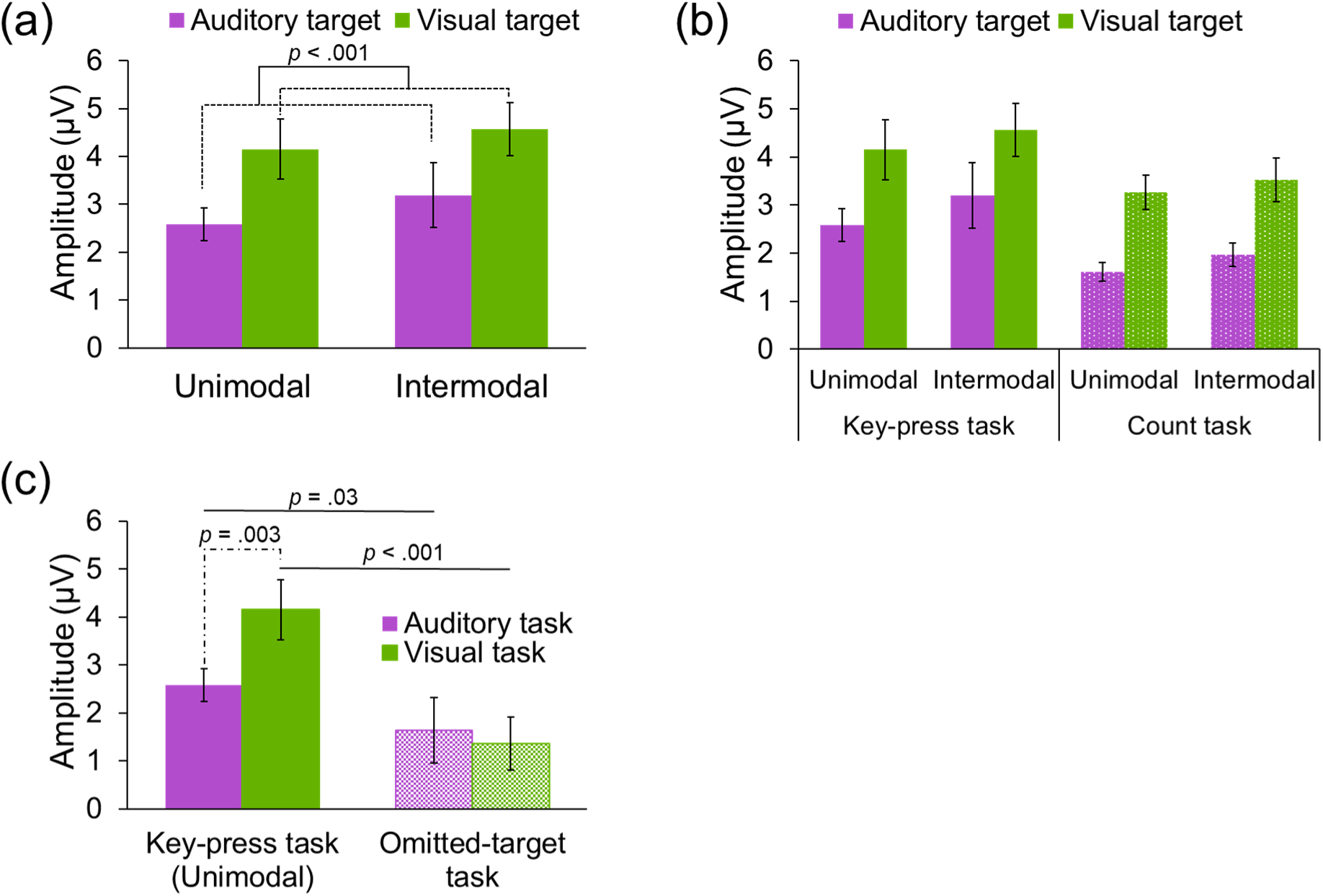
ERP results in the three experiments of this study. (a) P300 amplitude at the case of target modality and modality coupling in Experiment 1. (b) P300 differences between target modality, modality coupling, and response format in Experiment 2. (c) P300 differences between stimulus modality and the existence or absence of physical target stimuli in Experiment 3.

#### 3-2-3. Experiment 2: Examining the effects of the response format

Next, to investigate the effect of response format on P300 amplitude, a 2 (target modality: auditory, visual) × 2 (modality coupling: unimodal, intermodal) × 2 (response format: key press, count) mixed-factors ANOVA was conducted.

The results showed that the main effect of modality coupling (*F*(1,38) = 5.18, *p* = .03, η_p_ ^2^ = 0.12) and the main effect of target modality (*F*(1,38) = 55.87, *p* < .001, η_p_ ^2^ = 0.60) were significant as was the main effect of the response type (*F*(1,38) = 3.51, *p* = .07, η_p_ ^2^ = 0.08; Figure 5b). The P300 amplitude was larger when the intermodal condition was used rather than when the unimodal condition was used; the P300 amplitude was larger when the visual target was used than when the auditory target was used; and the P300 amplitude was larger when the participant responded to the target by pressing a key rather than when it was counted. None of the interactions were significant (*p*s > .57).

#### 3-2-4. Experiment 3: Examining the effects of the presence or absence of **physical target stimuli**

Finally, to investigate the effect of the presence or absence of a physical target stimulus on P300 amplitude, a 2 (stimulus modality: auditory, visual) x 2 (presence or absence of physical target: present, absent) mixed-factors ANOVA was conducted by comparing the unimodal auditory and visual data from Experiment 1 and the data from Experiment 3.

The results showed that the main effect of stimulus modality (*F*(1,37) = 13.57, *p* < .001, η_p_ ^2^ = 0.27) and the main effect of the presence or absence of a physical target (*F*(1,37) = 6.04, *p* = .02, η_p_ ^2^ = 0.14) were significant. In each case, the P300 amplitude was larger when the visual target was presented than when the auditory target was presented, and the P300 amplitude was larger when the physical target stimulus was presented than when it was not presented. Furthermore, the interaction was significant (*F*(1,37) = 12.50, *p* = .001, η_p_ ^2^ = 0.25). Post-hoc tests showed that the P300 amplitude was larger when there was a physical target stimulus than when there was no physical target stimulus, both in the auditory task (*F*(1,37) = 5.08, *p* = .03, η_p_ ^2^ = 0.12) and in the visual task (*F*(1,37) = 16.38, *p* < .001, η_p_ ^2^ = 0.31), and the P300 amplitude was larger for the visual task than for the auditory task only when there was a physical stimulus (*F*(1,19) = 11.22, *p* = .003, η_p_^2^ = 0.37; Figure 5c).

### 3-3. EEG microstate transition probabilities

Using the 10-state template with topographical polarity consideration (Figure 2) elucidated by Kashihara et al. (in press), we performed template fitting for the EEG epoch of each trial, each participant, each task, and each experiment to obtain the microstate label sequence. To focus on the whole-brain state transitions during the same period as the P300 component was identified in the ERP described in the previous section, template fitting was performed after sub-epoching from 300 to 600 ms following stimulus presentation.

The grand average transition probabilities (TPs) of the target (top row of each experiment) and standard trials (second row of each experiment) for each of the four tasks in all three experiments are shown in Figures 6—8. The TPs were calculated for 90 transitions, excluding self-recurrence, with the entire label sequence for a given epoch set to 1. Thereafter, the transition probabilities for each transition were averaged for each target trial (for a maximum of 80 trials) and standard trial (maximum of 320 trials) for each task within each participant and then averaged for each participant for each task to obtain the grand average TP for each task.

**Figure 6.**
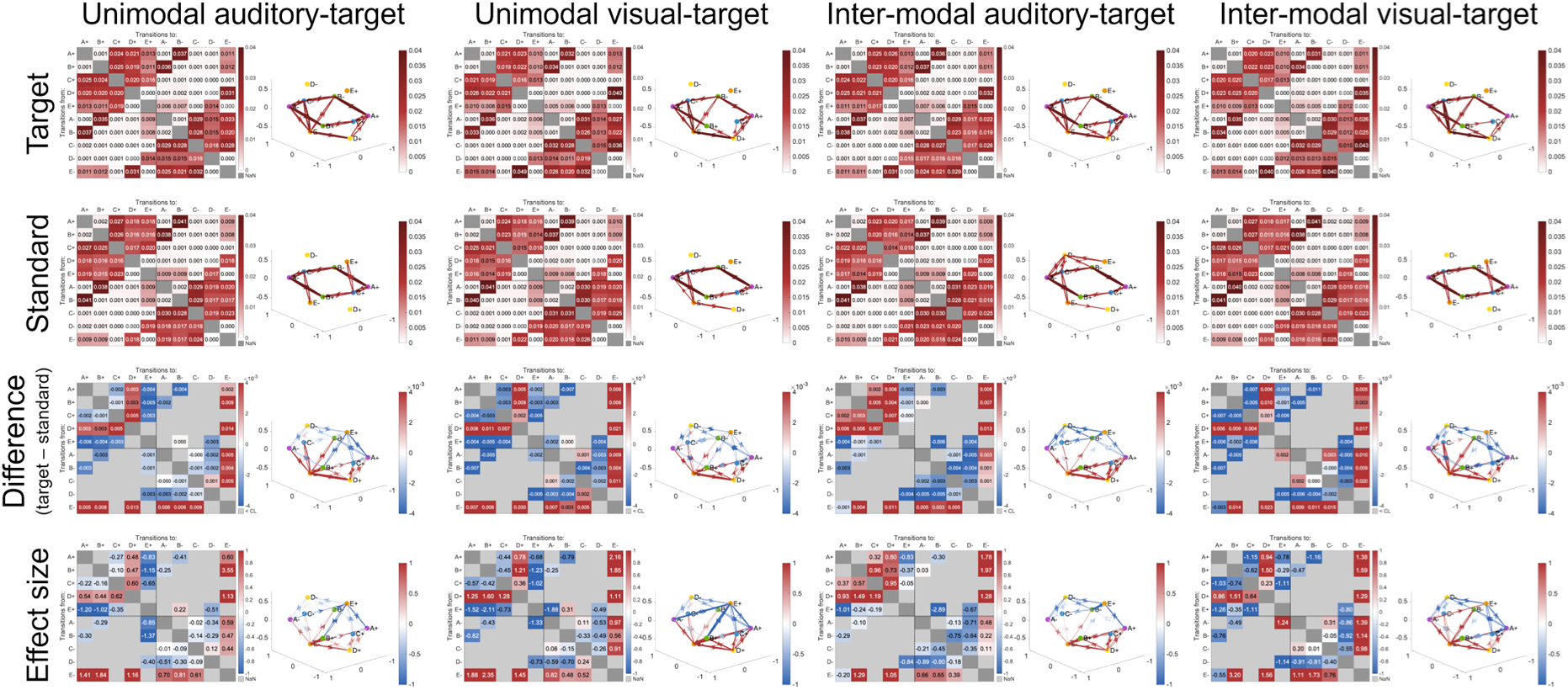
**EEG microstate transition probabilities in Experiment 1 of this study.**

**Figure 7.**
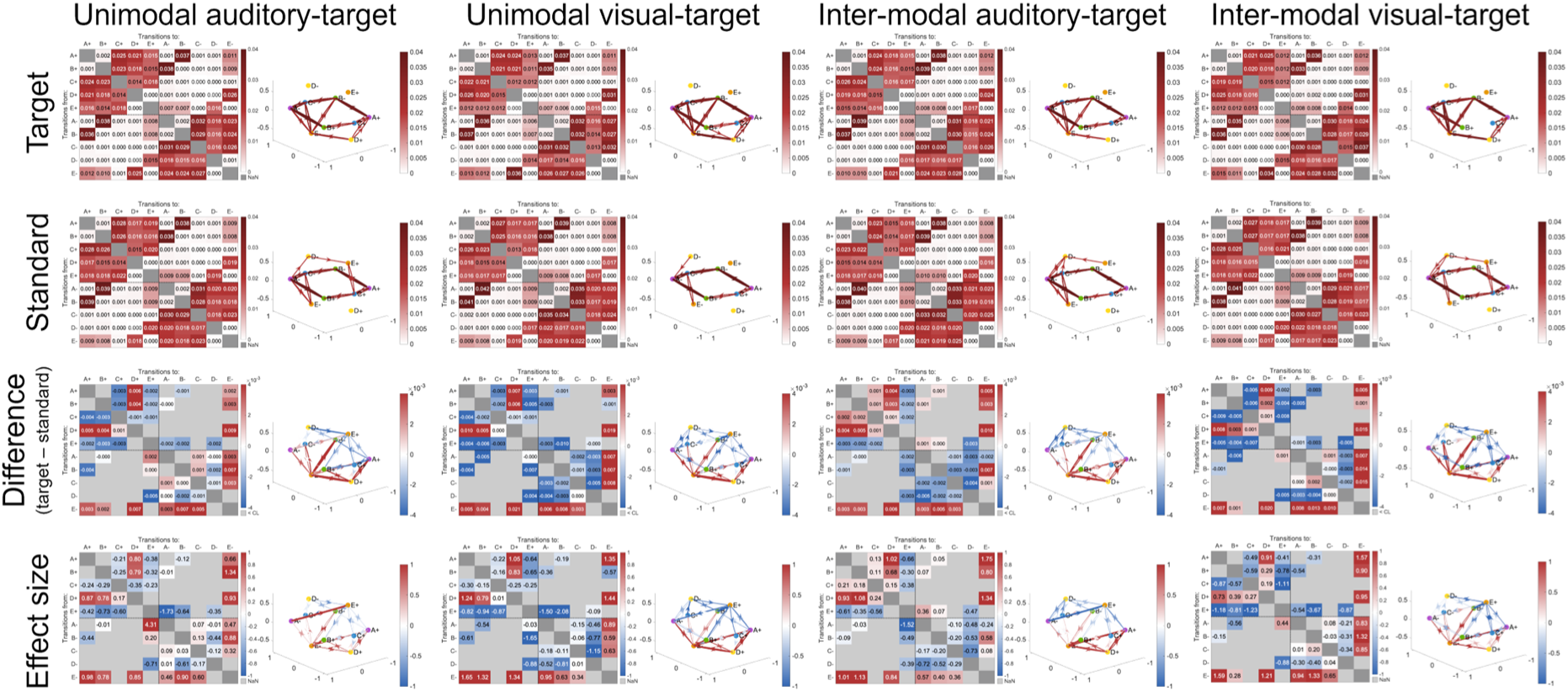
**EEG microstate transition probabilities in Experiment 2 of this study.**

**Figure 8.**
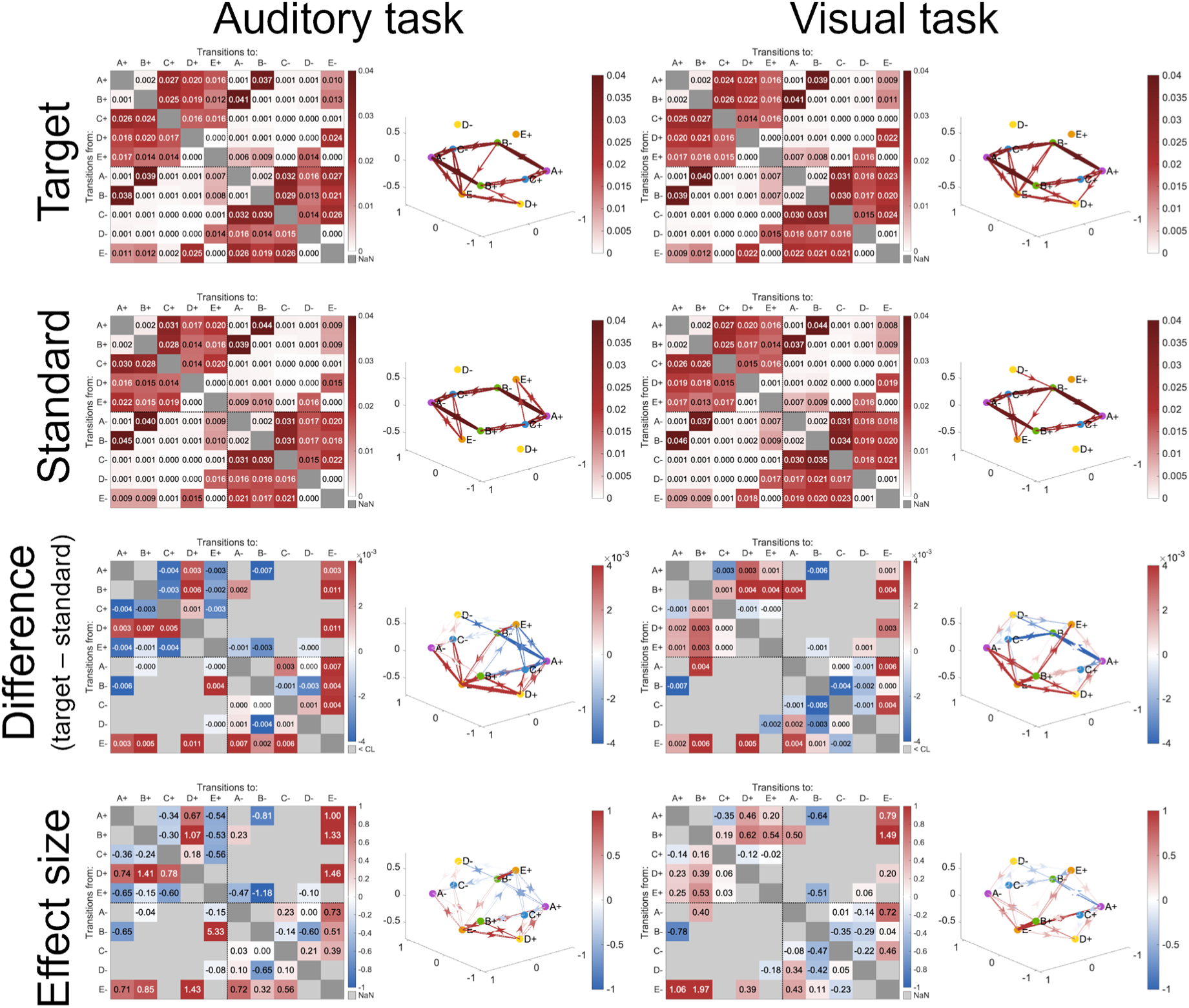
**EEG microstate transition probabilities in Experiment 3 of this study.**

To further confirm the difference between the TPs of the target and standard trials, we subtracted the grand average TP of the standard trials from that of the target trials to obtain the difference TP and standardized difference between the two (Figures 6—8, bottom two rows of each experiment). For the target and standard trials, similar trends were observed in all experiments and for all task conditions: compared to the standard trials, the target trials had more transitions that converged on E-between 300 and 600 ms after stimulus onset, and conversely, fewer transitions that related to the opposite pole, E+.

## 4. Discussion

In this study, we clarified the polarity-sensitive microstate transition dynamics in oddball tasks by conducting multiple oddball tasks with manipulated experimental conditions that are known to affect the P300 component. In Experiment 1, we examined the effects of the target stimulus modality and the coupling of the standard and target stimulus modalities on the EEG. Furthermore, to isolate the motor effects and the effects of the presence or absence of physical stimuli, we asked the participants to count without any motor behavior toward the target (Experiment 2), and we used a target without any physical stimulus (Experiment 3) and then compared these to Experiment 1.

In this study, very high response accuracy was obtained across all experiments and task conditions. In Experiments 1 and 2, there was no significant difference in the response accuracy based on the target stimulus modality and modality coupling. However, response times tended to be higher in the unimodal visual task. Furthermore, in Experiment 3, response times were higher and accuracy was lower in the visual task. These findings suggest that responding to visual stimuli may be more challenging than responding to auditory stimuli. Regarding response format, regardless of the stimulus condition coupling, accuracy tended to be higher when participants were required to press a key in response to the target, whether it was visual or auditory, rather than counting the occurrences. The difference between keypress responses, which involve processing the presented stimulus in real-time, and counting responses, which require the maintenance of a record of past occurrences while processing the current stimulus, may reflect differences in task goals and motivation among participants. The differences in response accuracy observed in this study may merely reflect the variations in task difficulty.

In this study, P300 ERP responses were observed in almost all oddball tasks. The amplitude of these P300 components was influenced by the target stimulus modality. Compared to when the target was an auditory stimulus, a target that comprised a visual stimulus elicited a higher amplitude P300. This finding is consistent with the results reported by Polich & Heine (1996), who showed that P300 from auditory stimuli was smaller than that from visual stimuli. Considering this finding alongside the behavioral results that suggest differences in task difficulty generates the inference that visual targets require a greater allocation of attentional resources than auditory targets, which are processed more passively. This increased attentional demand is reflected in the longer response times for visual stimuli and the corresponding P300 amplitude.

This hypothesis is supported by the results of Experiment 3, which investigated the effects of the presence or absence of a physical stimulus on the target. In Experiment 3, which focused on an unimodal task, the P300 amplitude was larger for the visual task than for the auditory task, but this was only observed when the physical target stimulus was presented. P300 could reflect brain activity during the modification of an environmental model, that is, during the updating of context (Donchin & Coles, 1988). Therefore, the effect of stimulus modality on P300 amplitude is an event that occurs depending on the characteristics of the physical target stimulus.

Regarding the effect of modality coupling between stimulus conditions on P300, Experiment 2 confirmed that, compared to the unimodal condition, the intermodal condition elicited higher P300 amplitudes. Although no significant difference was observed in Experiment 1, the descriptive trend was consistent. This finding aligns with previous studies (Brown et al., 2006, 2007) and extends prior research examining the influence of auditory and visual standard stimuli on late ERP components evoked by auditory targets. Brown et al. (2006, 2007) reported that late components, such as P300, are larger in audiovisual oddball tasks, which suggests the occurrence of distinct intermodal processes at later stages. Our study confirmed the occurrence of this late-stage context-dependent processing and suggests that it applies not only to auditory processing but also to visual processing.

Importantly, the target modality and the modality coupling between the target and standard stimuli each independently affected the P300 amplitude. This suggests that the meaning and context of each stimulus are important, regardless of whether the stimulus is visual or auditory or whether the frequent stimulus and rare target are the same or different modalities.

Furthermore, regarding the response format, the P300 amplitudes were greater in the keypress task than in the counting task. Prior research on oddball tasks has yielded inconsistent findings regarding the influence of response modality. However, our results support the notion that late ERP responses are stronger in keypress tasks.

We found that the ERP components were influenced by target stimulus modality, modality coupling, response format, and the presence or absence of physical stimuli. Previous ERP studies have suggested that these variables affect the configuration of EEG topography (e.g., Brown et al., 2006, 2007; Polich & Heine, 1996). To examine the spatial patterns of whole-brain states at a given moment and further track their temporal dynamics, polarity-informed microstate analysis was conducted to determine the transitional probabilities during the period corresponding to P300 generation. The results revealed that during the P300-related period, target trials exhibited increased cohesion toward a state referred to as E-. In contrast, the transitions associated with E+ tended to be less frequent than those in the standard trials. This finding suggests the process of transitioning to a specific state of a specific polarity rather than indication of an increase in a certain electrical field configuration pattern, and it also emphasizes the importance of considering topographical polarity in microstate analysis.

These transition patterns were consistently observed regardless of task conditions, response formats, or the presence or absence of physical stimuli. This suggests that the increased transition to E-may reflect a shift in whole-brain processing networks corresponding to the P300 component in target trials. The microstate E has been associated with interoceptive sensory and emotional information processing; moreover, its involvement in the salience network has been reported (Tarailis et al., 2024). The differentiation of the topographical polarity of microstates is a relatively recent development (Kashihara et al., in press), and little evidence has been accumulated regarding the functional distinctions between E+ and E-. However, fMRI studies on oddball tasks have suggested that key regions of the salience network, including the anterior cingulate cortex and insular cortex, are involved in this process (Harsay et al., 2012; Horn et al., 2003), indicating that E-may reflect the functions governed by the salience network. Specifically, the activation of the insular cortex, particularly the anterior insular cortex, potentially serves as an interface that mediates dynamic interactions between large-scale brain networks that are involved in externally directed attention and internally directed cognition (Menon & Uddin, 2010). That is, the salience network likely plays a critical role in detecting target stimuli, marking them for additional processing, and initiating appropriate control signals—such as keypress or counting responses. The spatiotemporal dynamics of EEG aggregation toward E-may reflect this process.

Finally, the limitations of this study should be acknowledged. First, this study manipulated experimental conditions to examine the effects of stimulus modality condition, response format, and the presence or absence of physical stimuli on relatively late neural dynamics following stimulus onset, which manifest as P300 and microstate transitions. However, the factors examined in this study represent only a subset of potential influences and other cognitive factors, such as attentional load and working memory demands, that may contribute to the formation of ERP components and microstate dynamics. Furthermore, investigating whole-brain state transitions using ERP markers in earlier processing stages may provide valuable insights. Future research should explore how these additional factors interact with the effects observed in the present study.

Moreover, through polarity-sensitive microstate analysis, we identified a potential role of transitions to E-in target stimulus detection and switching to appropriate processing states. However, the interpretation of polarity-sensitive microstate transitions remains an emerging area of research, and further studies are necessary to elucidate the distinct functional roles of E+ and E-within broader neural network dynamics. Although EEG microstate analysis offers high temporal resolution, its spatial resolution is inherently limited. Future research could integrate EEG with fMRI to more precisely localize the neural sources that mediate microstate transitions, particularly those associated with E-.

In conclusion, this study shows that EEG microstate transitions, in particular the increase in cohesiveness toward E-, correspond to the P300 component and reflect the dynamics of large-scale neural networks involved in target processing. This finding suggests that transitions to a whole-brain state pattern called E-play an important role in the detection and marking of target stimuli, facilitating the initiation of an appropriate response. Through the integration of microstate analysis with conventional ERP measurements, this study provided new insights into the spatiotemporal dynamics of neural network reconfiguration during the oddball task. These findings further the understanding of the neural mechanisms of stimulus-driven attention and highlight the importance of considering topographical polarity-sensitive microstate transitions in future research.

## Statements and Declarations

### Ethics approval

The study was approved by the Ethical Committee of ATR (approval numbers: 21-144 and 21-143).

### Availability of data and material

Data are available from the authors upon reasonable request.

## Acknowledgments

We are grateful to Yuki Inoue, Yoshiko Umemoto, and Ayako Oura for their assistance with data collection.

## Authors’ contributions

SK: Conceptualization, Methodology, Data curation, Investigation, Formal analysis, Visualization, Writing – original draft; TA: Conceptualization, Methodology, Validation, Writing – review and editing; HI: Project Administration, Supervision, Writing – review and editing. All authors read and approved the final manuscript.

